# Live-cell super-resolution nanoscopy reveals modulation of cristae dynamics in bioenergetically compromised mitochondria

**DOI:** 10.1101/2023.04.27.538553

**Authors:** Mathias Golombek, Thanos Tsigaras, Yulia Schaumkessel, Sebastian Hänsch, Stefanie Weidtkamp-Peters, Ruchika Anand, Andreas S. Reichert, Arun Kumar Kondadi

## Abstract

Cristae membranes have been recently shown to undergo intramitochondrial merging and splitting events. Yet, the metabolic and bioenergetic factors regulating them are unclear. Here we investigated whether and how cristae membrane remodelling is dependent on oxidative phosphorylation (OXPHOS) complexes, the mitochondrial membrane potential (ΔΨ_m_), and the ADP/ATP nucleotide translocator. Advanced live-cell STED nanoscopy combined with in-depth quantification were employed to analyse cristae morphology and dynamics after treatment of mammalian cells with rotenone, antimycin A, oligomycin A and CCCP. This led to formation of enlarged mitochondria along with reduced cristae density but did not change the number of cristae remodelling events. CCCP treatment leading to ΔΨ_m_ abrogation even enhanced the cristae dynamics showing their ΔΨ_m_-independent nature. Inhibition of OXPHOS complexes was accompanied by reduced ATP levels but did not affect cristae dynamics. However, inhibition of ADP/ATP exchange led to aberrant cristae morphology and impaired cristae dynamics in a mitochondrial subset. In sum, we provide quantitative data of cristae membrane remodelling under different conditions supporting an important interplay between OXPHOS, metabolite exchange and cristae membrane dynamics.

**Summary Blurb:** Cristae morphology and dynamics are intricately connected

## Introduction

Mitochondria are highly dynamic organelles playing vital roles in various cellular functions involving energy conversion, calcium buffering, iron-sulfur cluster biogenesis, immune responses, apoptosis and various metabolic reactions. Mitochondria consist of a smooth outer membrane (OM) and a heterogenous inner membrane (IM). Further, the IM is spatially divided into inner boundary membrane (IBM) and cristae membrane (CM). IBM runs parallel to the OM whereas CM invaginates towards the mitochondrial matrix. Crista junctions (CJs) are located at the interface between IBM and CM. CJs are slot- or pore-like structures typically having a diameter in the range of 25 nm (Frey & Mannella, 2000) which separate the intracristal space (ICS) and intermembrane space (IMS) adjacent to the IBM. CJs act as diffusion barrier for proteins (Davies *et al*, 2011; Gilkerson *et al*, 2003; Vogel *et al*, 2006; Wurm & Jakobs, 2006). In addition, CJs are proposed to play a role in the metabolite diffusion such as ATP, Ca^2+^, cytochrome *c* (Frey *et al*, 2002; Mannella *et al*, 2013). In fact, it has been shown that CJs establish electrochemical boundaries that allow differential membrane potential between the cristae and IBM (Wolf *et al*, 2019). In addition, individual cristae within a mitochondrion possess disparate membrane potentials demonstrating their functional independency. CJ formation was shown to depend on the high-molecular weight ‘Mitochondrial Contact Site and Cristae Organizing System’ (MICOS) complex consisting of at least seven proteins located at CJs (Harner *et al*, 2011; Hoppins *et al*, 2011; Rabl *et al*, 2009; von der Malsburg *et al*, 2011). A uniform nomenclature of MICOS subunits was established subsequently (Pfanner *et al*, 2014). MIC10, MIC13, MIC19, MIC25, MIC26, MIC27 and MIC60 constitute the proteins of the MICOS complex. The loss of CJs is observed upon depletion of most MICOS proteins leading to separation of IBM and CM (Harner *et al*., 2011; Hoppins *et al*., 2011; Stephan *et al*, 2020; von der Malsburg *et al*., 2011). MIC13 bridges the two subcomplexes: MIC60/19/25 and MIC10/26/27 (Anand *et al*, 2016; Guarani *et al*, 2015) via conserved WN and GxxxG MIC13 motifs (Urbach *et al*, 2021). Some subunits of the MICOS complex are evolutionarily conserved and have an endosymbiotic origin in α-proteobacteria (Eramo *et al*, 2020; Munoz-Gomez *et al*, 2023; Munoz-Gomez *et al*, 2015). The MICOS complex proteins are involved in mitochondrial protein import (von der Malsburg *et al*., 2011), lipid trafficking (Michaud *et al*, 2016), bending and remodeling the membranes (Hessenberger *et al*, 2017; Tarasenko *et al*, 2017) in addition to their role in the formation of CJs. Mutations of MICOS subunits have been associated with Parkinson’s disease, mitochondrial encephalopathy with liver disease and bilateral optic neuropathy, myopathy and lactic acidosis (Beninca *et al*, 2021; Guarani *et al*, 2016; Marco-Hernández *et al*, 2022; Tsai *et al*, 2018). Apart from the MICOS complex, F_1_F_O_ ATP synthase and OPA1 (Optic Atrophy Type I) play important roles in cristae remodelling as interplay of these complexes and cardiolipin is required for formation and maintenance of cristae and CJs (Anand *et al*, 2021; Kondadi *et al*, 2019, 2020b).

Cristae exists in various shapes and sizes depending on the cells, tissues and bioenergetic requirements (Zick *et al*, 2009). Moreover, alterations in cristae structure have been associated with neurodegeneration, diabetes, obesity, cardiomyopathy and myopathies (Eramo *et al*., 2020; Zick *et al*., 2009). For several decades, a static view of cristae prevailed based on the early existing electron microscopy (EM) data despite several indications of cristae remodeling. Cells undergoing apoptosis showed a massive reorganization of the mitochondrial IM where it reorganized and interconnected within a short span of few minutes (Scorrano *et al*, 2002). EM and electron tomography (ET) revealed that when isolated mitochondria were exposed to ADP, the IM confirmation changed to a highly interconnected network accompanied by matrix condensation termed State III respiration (Hackenbrock, 1966; Mannella *et al*, 1994; Perkins *et al*, 1997). Reduction in ADP levels resulted in a drastic decrease of interconnected cristae network accompanied by matrix expansion or state IV respiration. The application of SR techniques to resolve mitochondrial membranes has conclusively shown that cristae membranes are highly dynamic (Huang *et al*, 2018; Kondadi *et al*, 2020a; Kondadi *et al*., 2020b; Liu *et al*, 2022; Wang *et al*, 2019). Recently, we showed using live-cell stimulated emission depletion (STED) super-resolution (SR) nanoscopy that cristae membranes are dynamic and undergo continuous cycles of membrane remodeling dependent on the MICOS complex (Kondadi *et al*., 2020a). The cristae merging and splitting events are balanced, reversible and depend on the presence of the MICOS subunit MIC13. Fluctuation of membrane potential within individual cristae as well as photoactivation experiments at cristae-resolving resolution support the notion that cristae can exist transiently as isolated vesicles which are able to fuse and split with other cristae or the IBM (Kondadi *et al*., 2020a). It was recently shown that OPA1 and YME1L also affect cristae dynamics (Hu *et al*, 2020). However, it is unclear which metabolic factors are required to ensure cristae membrane dynamics, a process likely to consume considerably amounts of energy e.g. from ATP hydrolysis.

We asked what are the bioenergetic parameters which modulate the rates of cristae remodeling events: whether ATP levels or the mitochondrial membrane potential (ΔΨ_m_) influence cristae membrane dynamics and to which extent. In this endeavor, we performed advanced live-cell STED SR nanoscopy on mitochondria after inhibition of the electron transport chain (ETC) or the F_1_F_O_ ATP synthase using classical oxidative phosphorylation (OXPHOS) inhibitors. Furthermore, we used an OXPHOS uncoupler dissipating the ΔΨ_m_. Consistent with earlier studies using EM, we observed ∼50 % mitochondria to be enlarged, which showed decreased cristae density when compared to mitochondria which were not enlarged. We could further dissect and show that enlarged mitochondria in particular showed a moderate non-significant trend of increased cristae membrane dynamics. We conclude that the rate of cristae membrane dynamics is not negatively affected by inhibiting OXPHOS including dissipation of the ΔΨ_m_, reducing mitochondrial ATP levels but is rather moderately enhanced in enlarged mitochondria with reduced cristae density. This would be consistent with the view that cristae dynamics are either limited by structural constraints such as densely packed cristae or that reduction in cristae density is followed by an increased cristae fusion and fission rate as kind of a compensatory mechanism. Further, inhibition of adenine nucleotide translocator (ANT) by applying bongkrekic acid (BKA) to HeLa cells led to aberrant cristae morphology. Contrary to our observations using other OXPHOS inhibitors, we observed a clear reduction in cristae membrane dynamics in a subset of mitochondria, namely those showing aberrant cristae morphology. Overall, our results indicate that cristae membrane dynamics is linked to the bioenergetic state of mitochondria and point to a prominent role of the ADP/ATP metabolite exchange in this process.

## Results

### Cristae membrane dynamics is not impaired in mammalian cells treated with OXPHOS inhibitors

Recent application of novel SR techniques has revealed MICOS-dependent intramitochondrial cristae membrane dynamics in living cells (Kondadi *et al*., 2020a). However, the bioenergetic requirements which define such highly dynamic membrane remodeling processes are unknown. Here we utilized live-cell STED SR nanoscopy and determined dynamic alterations in cristae structure upon inhibition of ETC complexes, the F_1_F_O_ ATP synthase, or the ΔΨ_m_. For this, we treated HeLa cells with the following classical drugs: rotenone, antimycin A, oligomycin A inhibiting Complex I, Complex III, Complex V (F_1_F_O_ ATP synthase) respectively, and CCCP, a protonophore dissipating the ΔΨ_m_. We refer to these drugs collectively as mitochondrial toxins throughout the manuscript. Previously, we employed ATP5I-SNAP, marking F_1_F_O_ ATP synthase, to visualize cristae using live-cell STED SR nanoscopy in living cells (Kondadi *et al*., 2020a). Therefore, HeLa cells expressing ATP5I-SNAP were treated with silicon rhodamine (SiR) dye, which binds covalently to the SNAP-tag, followed by addition of mitochondrial toxins. SiR is suitable for SR imaging owing to its photostability and minimal fluorophore bleaching (Lukinavicius *et al*, 2013). We decided to follow cristae structure and cristae dynamics at very early time points, within 30 min, after addition of the respective toxins because of the following reasons: 1) To determine the immediate effect of acute bioenergetic alterations induced by different mitochondrial toxins on cristae remodeling. 2) To minimize secondary effects occurring later such as mitochondrial fragmentation which imposes methodological limitations for subsequent analyses. 3) Cells do not develop any obvious signs of cell death or undergo apoptosis for at least 1 hour after the addition of these mitochondrial toxins according to earlier reports (Duvezin-Caubet *et al*, 2006; Minamikawa *et al*, 1999). The concentrations of mitochondrial toxins used in this manuscript are broadly in the range used for real-time respirometry measurements where oxygen consumption has been shown to either increase or decrease depending on the mode of action of mitochondrial toxins used (Kondadi *et al*., 2020a; Stephan *et al*., 2020).

Using live-cell STED SR imaging, we followed cristae dynamics of cells treated with or without various mitochondrial toxins (**Fig 1A**). We achieved an image acquisition time of 0.94 s/frame which is improved compared to 1.2 to 2.5 s/frame achieved earlier (Kondadi *et al*., 2020a) using this technique. In all cases we observed that cristae showed robust remodeling events at a timescale of seconds independent of the presence of any mitochondrial toxin (**Fig 1A, Fig S1 and Videos 1-5**). Within the time span of each movie, we observed that cristae remodeling constantly showed the formation and reshaping of X- and Y-like structures observed in our previous study (Kondadi *et al*., 2020a). There was no apparent difference in the way cristae appeared or remodeled between the different toxins. Yet to analyze this in more detail and to test whether we missed subtle changes, we performed a blind quantification of cristae merging and splitting events within individual mitochondria. In our previous study, we obtained a spatial resolution of 50-60 nm using live-cell STED nanoscopy (Kondadi *et al*., 2020a) meaning that cristae with a distance more than 60 nm between them can be distinguished. We found that treatment of cells with mitochondrial toxins did not lead to any significant changes in the frequency of merging or splitting events in mitochondria (**Fig 1B**). We conclude that inhibition of mitochondrial OXPHOS complexes by these toxins does not impair cristae dynamics or cause an imbalance of merging and splitting events.

**Figure 1.**
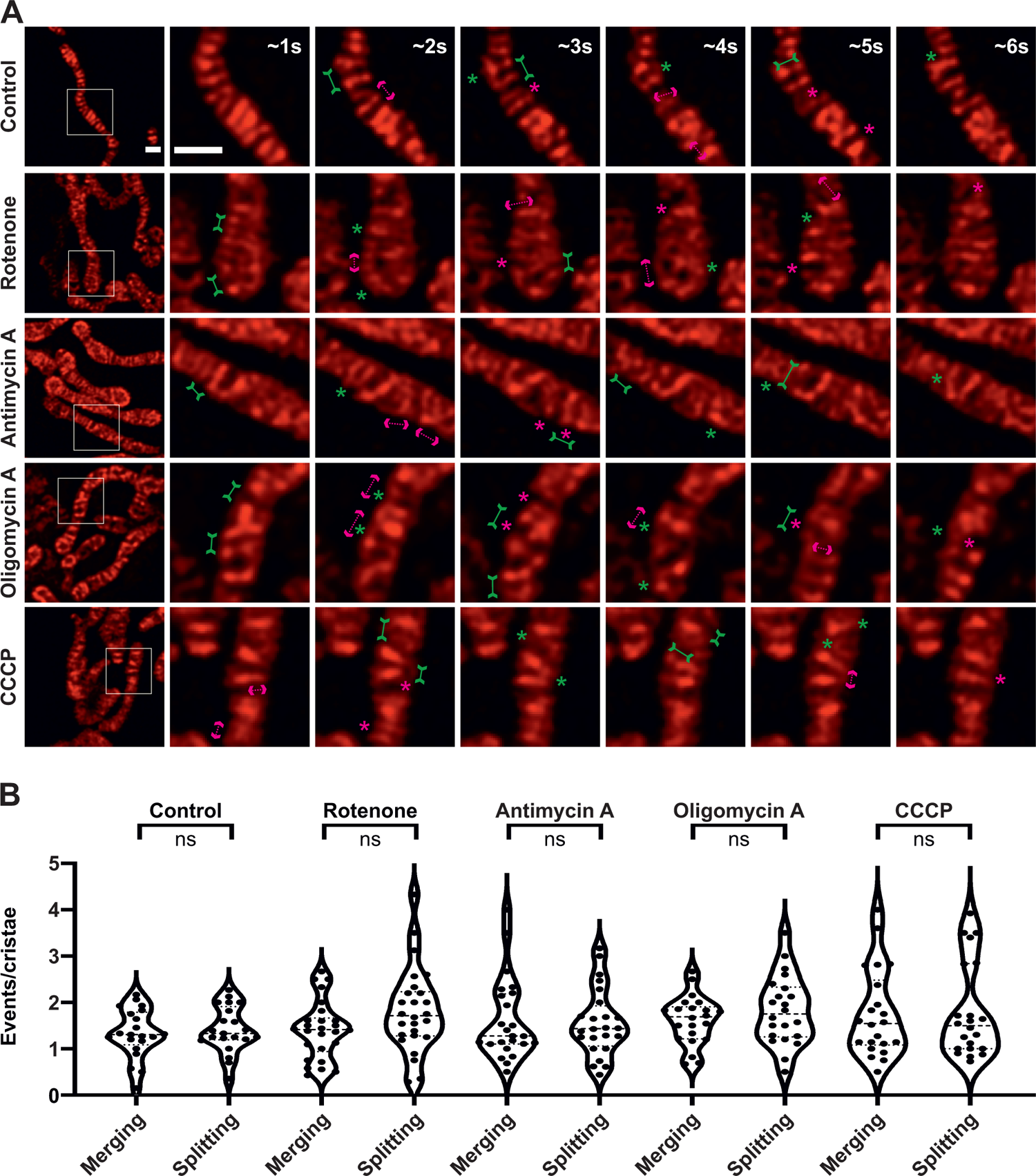
Crista merging and splitting events occur in a balanced manner upon inhibition of OXPHOS complexes and ΔΨ_m_ dissipation. **(A)** Representative live-cell STED SR images of HeLa cells, expressing ATP5I-SNAP and stained with silicon rhodamine, untreated or treated with rotenone, antimycin A, oligomycin A and CCCP. Images at the extreme left show whole mitochondria along with white inset boxes. Other images on the right-side display time-lapse series (0.94 s/frame) of zoom of mitochondrial portion in the white inset at ∼1 s, 2 s, 3 s, 4 s, 5 s and 6 s. Green and magenta asterisks show corresponding merging and splitting events while solid green arrows pointing inward and dotted magenta arrows pointing outward show imminent merging and splitting events respectively. Scale bar represents 500 nm. **(B)** Blind quantification of cristae merging and splitting events per mitochondrion in different conditions as described in **(A)**. Pooled data from three individual experiments with 21 to 26 mitochondria is shown as violin plots with individual data points. Each symbol represents one mitochondrion. (*ns = nonsignificant P-value* > 0.05). In order to increase the number of cells considered for quantification, a maximum of two mitochondria were considered from a single cell, throughout the manuscript, where STED nanoscopy was performed. One-way ANOVA was used for statistical analysis.

### Cristae structure is altered in a subset of mammalian cells treated with mitochondrial toxins

To further analyze whether alterations in cristae dynamics is eventually linked only to a subset of mitochondria, we performed a detailed characterization of cristae architecture upon treatment with various toxins. We noted that the treatments of HeLa cells with various mitochondrial toxins led to the formation of mitochondria with increased width in a fraction of cells (**Fig 2A,** bottom panel), while another fraction did not show a change in mitochondrial width (**Fig 2A,** top panel) and resembled control cells. We categorized percentage of mitochondria possessing corresponding mitochondrial widths under each condition **(Fig 2B-F)**. Frequency distribution curves revealed that control cells displayed a gaussian-like distribution of mitochondrial width with the highest percentage of mitochondria present in 400-500 nm range while a maximum mitochondrial width of 600 nm was observed (**Fig 2B**). On the contrary, treatment of cells with mitochondrial toxins led to substantially increased mitochondrial width (**Fig 2A,** bottom panel **and 2C-F**) as shown by shift towards the right in percentage mitochondria. We found that irrespective of the toxin used around 50 % of the mitochondria (maximum two mitochondria considered per cell) were enlarged (width ≥ 650 nm), and that no mitochondria under control conditions had an average width larger than or equal to 650 nm (**Fig 2B-F**). Hence, mitochondrial dysfunction induced by rotenone, antimycin A, oligomycin A and CCCP uniformly led to enlarged mitochondria within 30 min. Based on these results, we used 650 nm mitochondrial width as the cut-off for defining mitochondria as ‘enlarged’ (from here on) as this excluded all the mitochondria from the control group (referred to as ‘normal’ mitochondria here on). We next quantified cristae structure-related parameters for all mitochondria including distributing them into subsets of normal and enlarged mitochondria. We characterized cristae number per µm^2^ of mitochondrial area defined as cristae density, average distance between cristae in a mitochondrion defined as intercristae distance and the percentage area occupied by cristae within a mitochondrion. We did not find major differences in different cristae parameters described above when we compared the entire population of mitochondria in cells treated with or without various mitochondrial toxins **(Fig 2G, Fig S2A, C)**. Still, we observed an apparent trend indicating that cristae density is negatively correlated with mitochondrial width when mitochondrial toxins were applied **(Fig S2E-I)**. Control HeLa cells exhibited a median cristae density of around 7 cristae/µm^2^ which was similar to cells treated with mitochondrial toxins (**Fig 2G**). When we distributed the mitochondria as having normal or enlarged width, we found that mitochondria with normal mitochondrial width showed similar cristae density compared to untreated mitochondria (**Fig 2H**). Enlarged mitochondria exposed to mitochondrial toxins had a median cristae density of 4 cristae/µm^2^ compared to 7 cristae/µm^2^ in control cells. Only mitochondria showing enlarged width showed a statistically significant decrease of the cristae density for all toxins when compared to control mitochondria (**Fig 2H**). These findings are well recapitulated by the observed increased trend in the average intercristae distance which is altered again for enlarged mitochondria (**Fig S2 A, B**). We next checked whether applying mitochondrial toxins led to a change in the percentage cristae area occupied per mitochondrion. We observed that the percentage cristae area per mitochondrion was unchanged upon addition of mitochondrial toxins within the time window of imaging (**Fig S2C**) which was independent of the mitochondrial width (**Fig S2D**). Taken together, STED SR nanoscopy revealed that bioenergetically compromised mammalian cells within a short time span result in structural changes where ∼50% of mitochondria are characterized by decreased cristae density, increased average distances between adjacent cristae with no gross changes in relative cristae area occupied by mitochondria. These observations are reflected in the negative correlation of cristae density and mitochondrial width **(Fig S2E-I)**. We next aimed to check ultrastructural changes under these conditions using EM. Consistent with results obtained using STED nanoscopy (**Fig 2A-F & Fig S2**), electron micrographs revealed enlarged mitochondria and increased distance between the cristae (shown using white arrows) upon treatment of HeLa cells with all mitochondrial toxins (**Fig 2I**). Increased distances between the cristae contributed to a visible decrease in the cristae density compared to control mitochondria which was in line with previous observations that cells treated with different mitochondrial toxins resulted in enlarged mitochondria accompanied by decreased cristae density (Gottlieb *et al*, 2003; Hytti *et al*, 2019). Overall, using a combination of STED SR nanoscopy and EM, we show that treatment of HeLa cells with various mitochondrial toxins, which disrupt the ETC function and ΔΨ_m_, resulted in enlarged mitochondria accompanied by increased intercristae distance and reduced cristae density.

**Figure 2.**
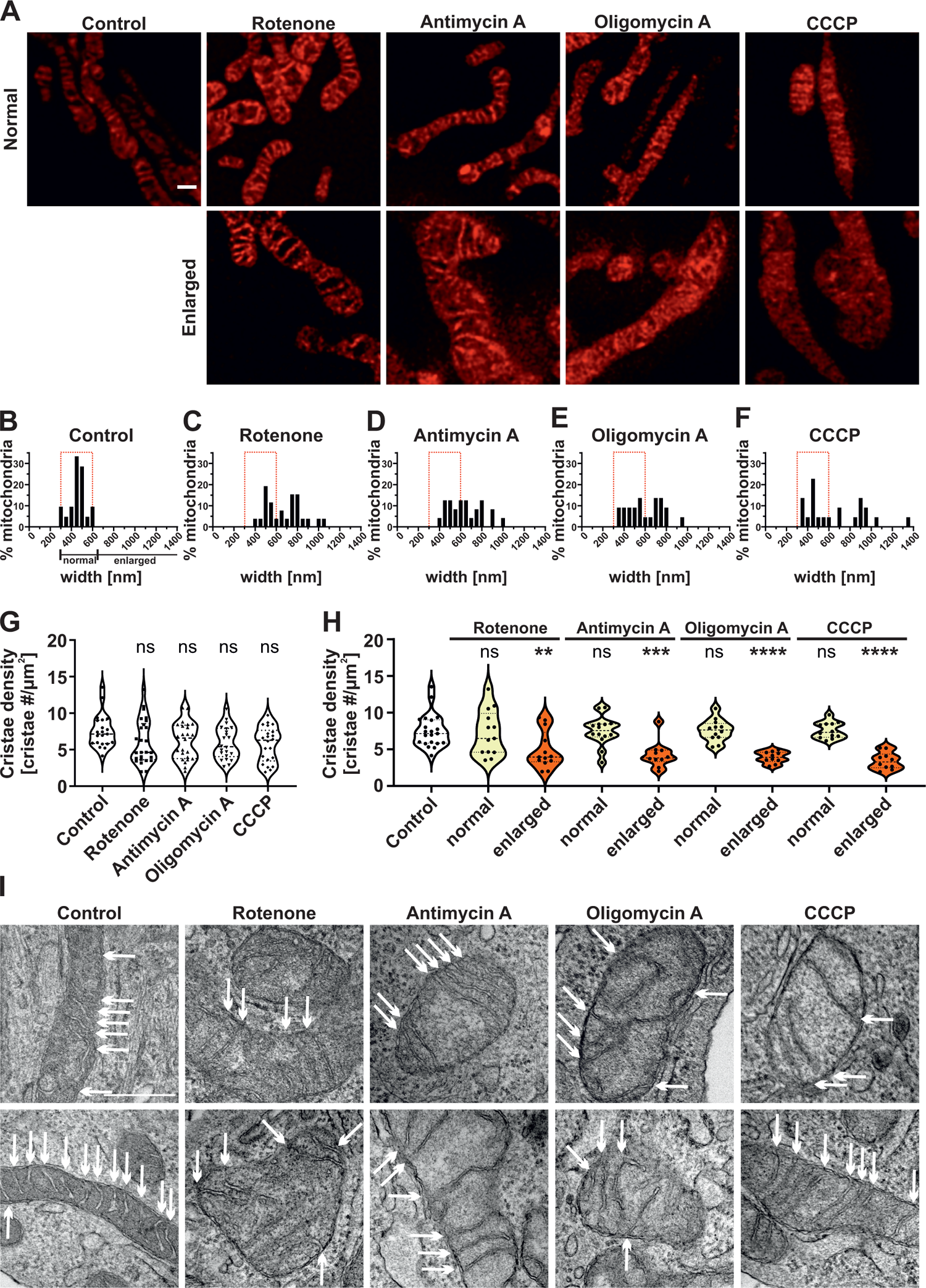
Mitochondrial toxins alter the morphology of cristae as well as mitochondria. **(A)** Representative STED SR images of HeLa cells expressing ATP5I-SNAP, stained with silicon rhodamine, displaying normal (< 650 nm) or enlarged (≥ 650 nm) mitochondrial width upon rotenone, antimycin A, oligomycin A and CCCP treatment. Top and bottom rows show mitochondria with normal and enlarged width respectively. Scale bar represents 500 nm. **(B-F)** Frequency distribution (50 nm bins) of percentage mitochondria having particular mitochondrial width in control cells **(B)** and cells treated with rotenone **(C)**, antimycin A **(D)**, oligomycin A **(E)** and CCCP **(F)** obtained from three independent experiments (21 to 26 mitochondria). Red rectangle indicates width distribution of untreated control group which was superimposed in toxin-treated conditions. **(G-H)** Quantification of cristae density (cristae number per mitochondrial area in µm^2^) per mitochondria, **(G)** Pooled data from three individual experiments is shown as violin plots with individual data points (21 to 26 mitochondria). Each symbol represents one mitochondrion. One-way ANOVA was used for statistical analysis. **(H)** Data was separated into normal and enlarged based on mitochondrial width with each condition having 10 to 21 mitochondria. Conditions were compared to untreated control group. (*ns = nonsignificant P-value* > 0.05, *\*\*P-value* ≤ 0.01, *\*\*\*P-value* ≤ 0.001, *\*\*\*\* P-value* ≤ 0.0001). One-way ANOVA was used for statistical analysis. **(I)** Representative transmission electron micrographs of mitochondria of cells treated without or with rotenone, antimycin A, oligomycin A and CCCP. Individual cristae within a mitochondrion are marked using white arrows. A higher number of arrows in control mitochondria show increased number of cristae per mitochondrial section when compared to mitochondria where the cells were treated with various mitochondrial toxins. Two mitochondria are shown per condition. Scale bar represents 500 nm.

### Cristae membrane dynamics is unchanged in enlarged mitochondria treated with various mitochondrial toxins

Given the structural alterations in a subset of mitochondria and the finding that cristae dynamics is overall robustly occurring in bioenergetically compromised mitochondria (**Fig 1**), we wondered whether cristae dynamics is specifically altered in mitochondria that have been structurally altered and the overall effect was masked. Upon treatment with mitochondrial toxins, in enlarged mitochondria, the frequencies of merging and splitting events remained balanced and we observed X- and Y-like structures appearing and disappearing at a timescale of seconds **(Fig 3A, S3A-C and videos 6-10)**. Overall, when we re-visited normal and enlarged mitochondria separately, we observed an apparent, yet no significant increase in the frequency of both merging and splitting events in enlarged mitochondria after cells were treated with antimycin A and oligomycin A but not in normal mitochondria after the same treatments (**Fig 3A-E)**. However, in enlarged mitochondria a significant increase of splitting events was observed after rotenone treatment as well as of merging and splitting events when the ΔΨ_m_ was dissipated by CCCP **(Fig 3C, E)**. Also, we rule out that cristae membrane dynamics is reduced upon inhibition of OXPHOS in the subset of enlarged mitochondria. In contrast, cristae membrane dynamics is moderately increased after dissipating the ΔΨ_m_.

**Figure 3.**
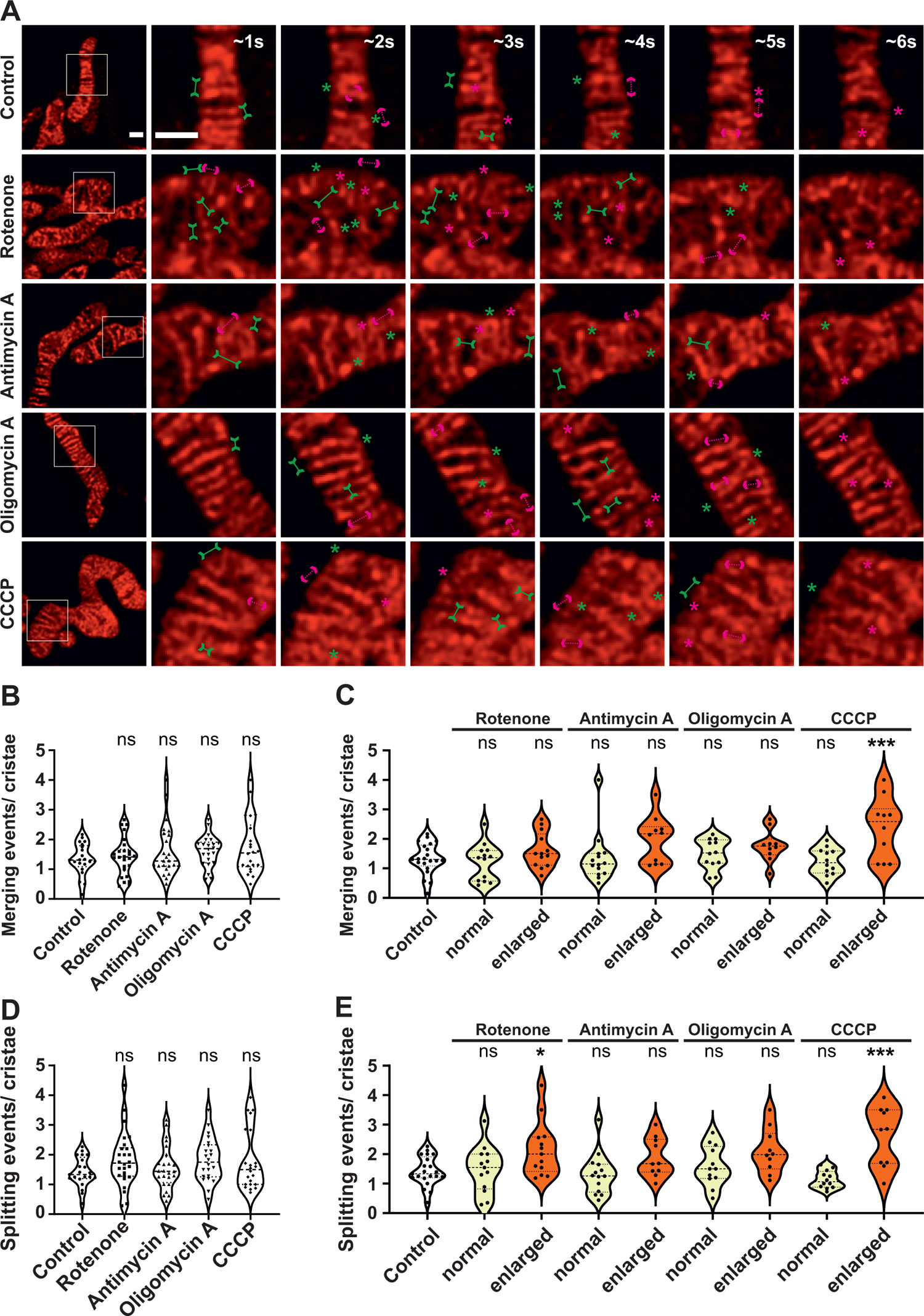
Crista merging and splitting events are maintained in enlarged mitochondria. **(A)** Representative live-cell STED SR images of HeLa cells, expressing ATP5I-SNAP and stained with silicon rhodamine, showing control and enlarged mitochondria obtained after treatment without or with various mitochondrial toxins respectively. Images at the extreme left show whole mitochondria along with white inset boxes. Other images on the right-side display time-lapse series (0.94 s/frame) of zoom of mitochondrial portion at ∼1 s, 2 s, 3 s, 4 s, 5 s and 6 s. Green and magenta asterisks show corresponding merging and splitting events while solid green arrows pointing inward and dotted magenta arrows pointing outward show imminent merging and splitting events respectively. Scale bar represents 500 nm. **(B-E)** Blind quantification of cristae merging and splitting events per mitochondrion in different conditions described in **(A)**. **(B)** Quantification of cristae merging events per mitochondrion from three individual experiments (21 to 26 mitochondria) is shown as violin plots with individual data points. Each symbol represents one mitochondrion. **(C)** The number of cristae merging events were classified into normal (< 650 nm) or enlarged (≥ 650 nm) mitochondria, with each condition having 10 to 21 mitochondria. Mitochondrial toxin treatment conditions were compared to untreated control group. **(D)** Quantification of cristae splitting events per mitochondrion from three individual experiments (21 to 26 mitochondria) is shown as violin plots with individual data points. Each symbol represents one mitochondrion. **(E)** The number of cristae splitting events were classified into normal (< 650 nm) or enlarged (≥ 650 nm) mitochondria, with each condition having 10 to 21 mitochondria. Different conditions were compared to untreated control group. (*ns = nonsignificant P-value* > 0.05, *\* P-value* ≤ 0.05, *\*\*\*P-value* ≤ 0.001). One-way ANOVA was used for statistical analysis.

In order to check how the mitochondrial ATP levels were influenced by various mitochondrial toxins and whether there was a correlation between cristae membrane dynamics and ATP levels, we checked the mitochondrial ATP levels after the respective treatments within the time span of 30 min similar to STED nanoscopy. For determining mitochondrial ATP levels, we took advantage of the mitGO-Ateam2 probe (Nakano *et al*, 2011). mitGoTeam2 is a ratiometric intramolecular FRET probe which binds ATP to bring the GFP, acting as FRET donor, close to OFP, the FRET acceptor, leading to an increased acceptor emission. Hence, reduction of ATP levels leads to decrease in the ratio of emission maximum at 580 nm (OFP)/520 (GFP) nm. mitGO-Ateam2 and mitoAT1.03 probes have been used to study spatiotemporal modulations of mitochondrial ATP levels (Imamura *et al*, 2009; Nakano *et al*., 2011). Pseudocolor ratiometric rainbow LUT images clearly showed that emission of 580 nm/520 nm significantly decreased in mitochondria of cells treated with rotenone, antimycin A and oligomycin A when compared to control HeLa cells (**Fig 4A,** bottommost panel). Accordingly, quantification of the ratio of emission maximum at 580 nm/520 nm showed that all cells treated with rotenone, antimycin A and oligomycin A displayed a significant reduction of ATP levels (**Fig 4B**). Surprisingly, treatment of HeLa cells with CCCP did not show any change in the ATP levels within the short time span of 30 min. Further, we checked whether the reduction of ATP levels was affected by mitochondrial width. For this, we compared the ATP levels of normal and enlarged mitochondria in cells treated with OXPHOS inhibitors but did not find any differences (**Fig 4C**). This suggests that the observed reduction in mitochondrial ATP levels in cells treated with mitochondrial toxins precedes formation of enlarged mitochondria. Overall, we conclude that unaltered cristae dynamics in enlarged mitochondria is not due to delayed or inefficient action of mitochondrial toxins demonstrating that cristae dynamics is robustly maintained at reduced ATP levels.

**Figure 4.**
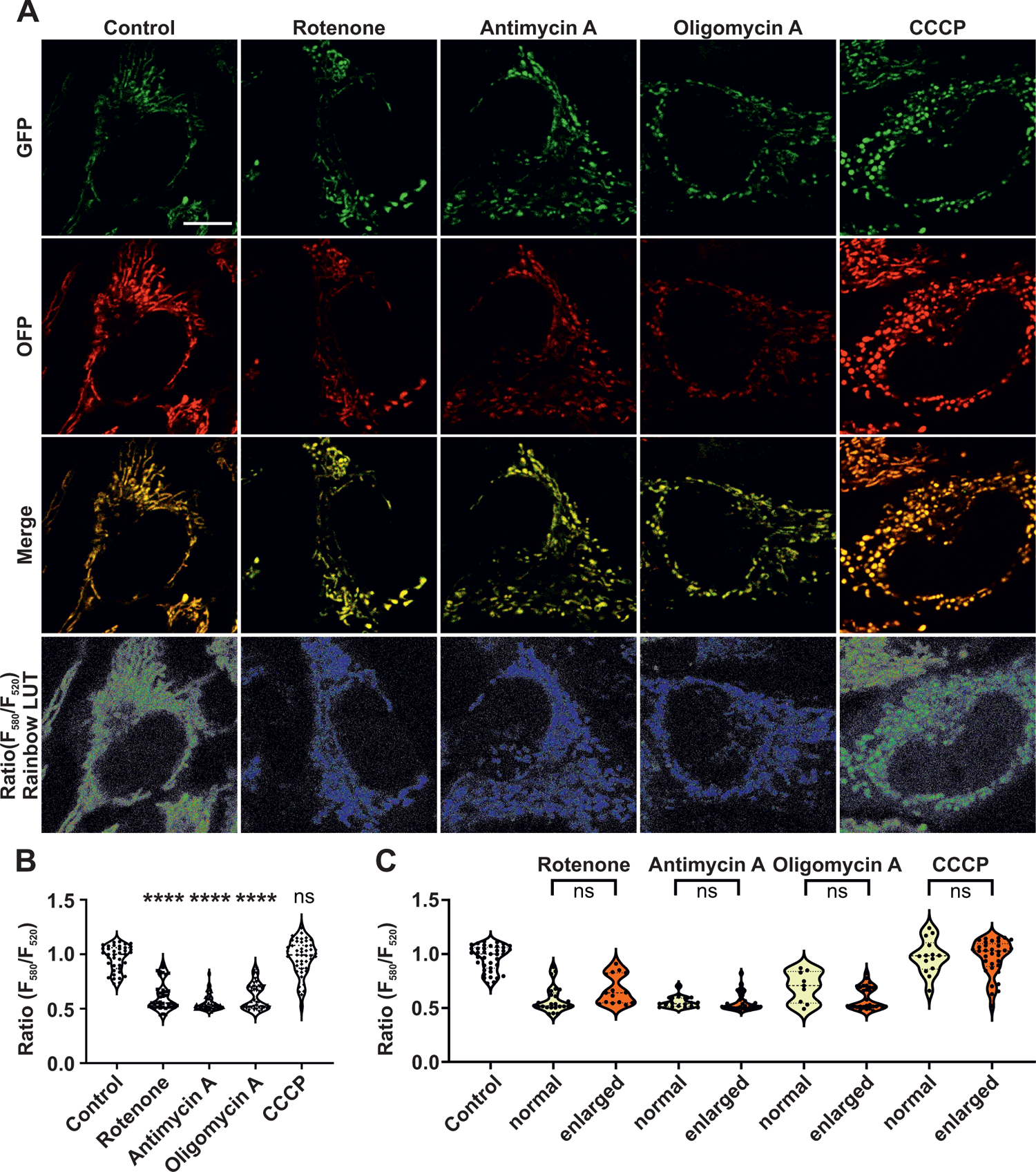
Mitochondrial ATP levels are significantly reduced upon inhibition of ETC complexes I, III and V. **(A)** Representative images of HeLa cells expressing mitGOTeam2, a ratiometric FRET-based genetically-encoded sensor determining the ATP levels, in cells treated without or with rotenone, antimycin A, oligomycin A and CCCP. The images in first row show the FRET donor emission (GFP) while the images in second row display the FRET acceptor emission (OFP). The third row represents a merge of FRET donor and acceptor emission channels. The bottommost row represents ratiometric 32-bit float images, shown using pseudocolour rainbow LUT intensities, used as a basis for quantifying mitochondrial ATP levels. Rainbow LUT intensities reveal low-intensity blue pixels in cells exposed to mitochondrial toxins compared to high-intensity green and red pixels in untreated control cells. Scale bar represents 10 µm. **(B-C)** Quantification of cellular mitochondrial ATP levels obtained by dividing the intensities of FRET acceptor emission (580 nm) by FRET donor emission (520 nm) in HeLa cells treated with or without the mentioned mitochondrial toxins. **(B)** Quantification of mitochondrial ATP levels (ratiometric data) is shown as violin plots from three individual experiments (39 to 50 cells). Each symbol represents mitochondrial ATP levels of an individual cell. Conditions are compared to untreated control group. **(C)** Ratiometric data was separated into cells with either prevalent normal or enlarged mitochondria (described in methods). Cells with mixed mitochondrial morphology were excluded from this evaluation resulting in 9 to 39 cells for each group. Statistical analysis was preformed between the two classified groups for each treatment condition, with untreated control group as reference. (*ns = nonsignificant P-value* > 0.05, *\*\*\*\* P-value* ≤ 0.0001). One-way ANOVA was used for statistical analysis.

Next, we addressed how ΔΨ_m_ is influenced after applying the respective mitochondrial toxins in the time window which was used for STED SR nanoscopy and ratiometric FRET-based ATP level detection. Antimycin A and CCCP strongly decreased ΔΨ_m_ (**Fig 5A**). Accordingly, detailed quantification revealed a significant decrease of ΔΨ_m_ when cells were treated either with antimycin A or CCCP when compared to untreated mitochondria (**Fig 5B**). There was a modest but significant decrease of ΔΨ_m_ with rotenone treatment while cells treated with oligomycin showed no change in the ΔΨ_m_. When we put the rate of cristae dynamics (**Fig 3**) in the context of ΔΨ_m_ measurements (**Fig 5**), we find that there was no reduction in the rate of cristae merging and splitting events in enlarged mitochondria upon introduction of rotenone, antimycin A and CCCP compared to control mitochondria despite a significant decrease of ΔΨ_m_. On the contrary, cells treated with F_1_F_O_ ATP synthase inhibitor, oligomycin A, exhibited no significant change in the number of merging and splitting events compared to control mitochondria but did not show a decrease of ΔΨ_m_ which was still accompanied by a significant reduction in ATP levels. Overall, we conclude that the merging and splitting events occur in enlarged mitochondria independent of ΔΨ_m_. Thus, cristae dynamics appear to operate independent of the ΔΨ_m_ and are maintained even at reduced ATP levels.

**Figure 5.**
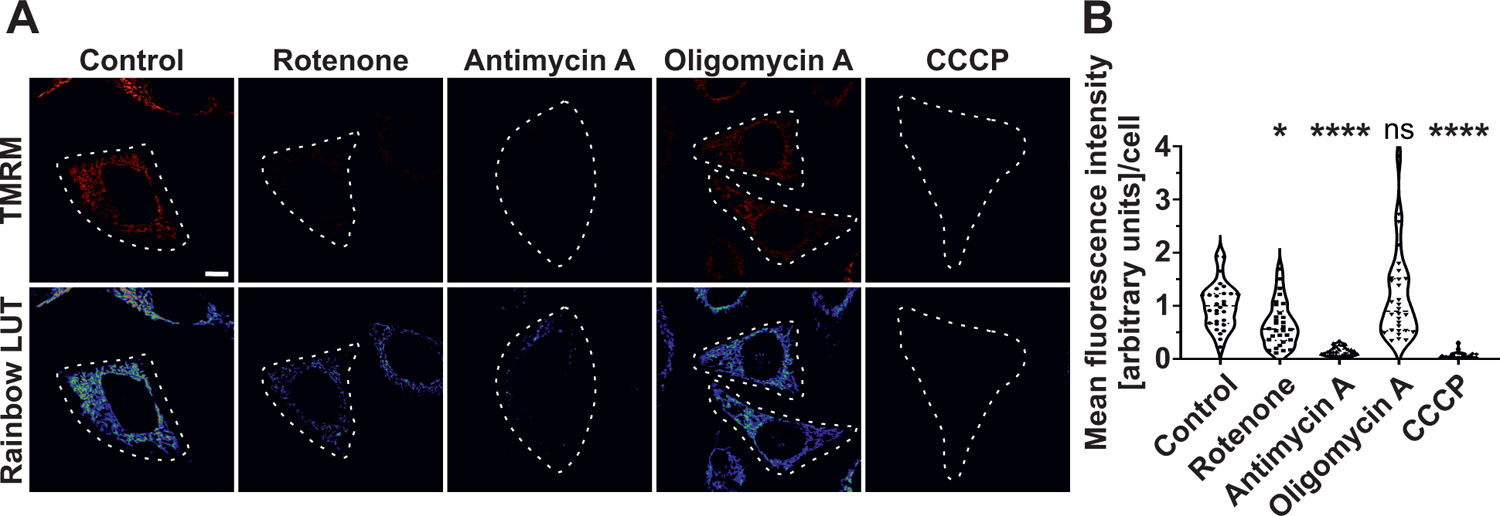
Mitochondrial membrane potential is depleted upon treatment with rotenone, antimycin A and CCCP but not oligomycin A. **(A)** Representative confocal images of HeLa cells (indicated by white dotted line) stained with TMRM either untreated or treated with rotenone, antimycin A, oligomycin A and CCCP (top row). Corresponding pseudocolour rainbow LUT intensities of respective TMRM signal is shown in bottom row. Scale bar represents 10 µm. **(B)** Quantification of ΔΨ_m_ based on mean TMRM fluorescence intensity measurements of individual HeLa cells either treated without or with various mitochondrial toxins mentioned. Results are shown as violin plots, with all individual data points. Data are obtained from 3 independent experiments (52 to 65 cells). Statistical comparisons were drawn between the untreated control group and the toxin-treated conditions. (*ns = nonsignificant P-value* > 0.05, *\* P-value* ≤ 0.05, *\*\*\*\* P-value* ≤ 0.0001). One-way ANOVA was used for statistical analysis.

### Cristae morphology is perturbed when HeLa cells are treated with an inhibitor of the adenine nucleotide translocator (ANT)

We next asked whether ADP/ATP exchange of mitochondria, mediated by the ANT, is regulating the dynamics of cristae membranes. In this context, we used various concentrations of bongkrekic acid (BKA), an ANT inhibitor, at 10 µM, 25 µM and 50 µM on HeLa cells. A clear dose-dependent decrease of oxygen consumption was observed with increasing concentration of BKA within the time window (30 min) used for imaging cristae membrane dynamics and ATP levels throughout this manuscript (**Fig 6A, B**). We used the highest concentration (50 µM) of BKA for imaging cristae morphology and dynamics as it showed the strongest decrease in mitochondrial oxygen consumption. We noted that cristae morphology was clearly aberrant upon BKA treatment compared to untreated controls (**Fig 6C, D**). STED nanoscopy revealed numerous mitochondria with huge spaces devoid of cristae as well as highly interconnected cristae upon BKA treatment (**Fig 6C, D,** top panel, **E and Fig S4A, B**) and also other abnormal cristae organization where cristae were either clumped or accumulated in the central region of swollen mitochondria (**Fig S4C**). Overall, the alterations in cristae morphology observed by STED imaging were validated when EM was employed (**Fig 6C, D,** bottom panel). Therefore, we characterized the percentage of mitochondria having normal and abnormal cristae morphology. There was a clear increase of mitochondria which had abnormal cristae morphology in BKA-treated condition when compared to untreated cells when the data from all five experiments were pooled as ∼33 % mitochondria had abnormal cristae morphology when compared to ∼13 % mitochondria in untreated conditions (**Fig S4A, B**). There were instances where live-cell STED movies of BKA-treated mitochondria showed highly interconnected cristae where the cristae dynamics were apparently highly reduced or static (**Fig 6E and videos 11-13**). We quantified the cristae dynamics in control as well as BKA-treated mitochondria and found no change in the overall merging and splitting events (**Fig S4D**). However, BKA-treated mitochondria with abnormal cristae morphology showed significantly reduced cristae dynamics compared to all mitochondria with BKA treatment or untreated mitochondria (**Fig 6F, G**). The cristae merging and splitting events were still balanced in control cells and BKA-treated cells (**Fig S4 D**). The ΔΨ_m_ was significantly decreased in BKA-treated cells (**Fig S4E, F**). Thus, despite the overall decrease in ΔΨ_m_, a reduction in cristae dynamics was observed only in those mitochondria where the cristae morphology was aberrant. Overall, we conclude that inhibition of the ANT by BKA results in alteration of cristae morphology and a partial reduction of cristae membrane dynamics suggesting that ADP/ATP exchange across the inner membrane is critical to maintain to cristae membrane dynamics independent of the membrane potential.

**Figure 6.**
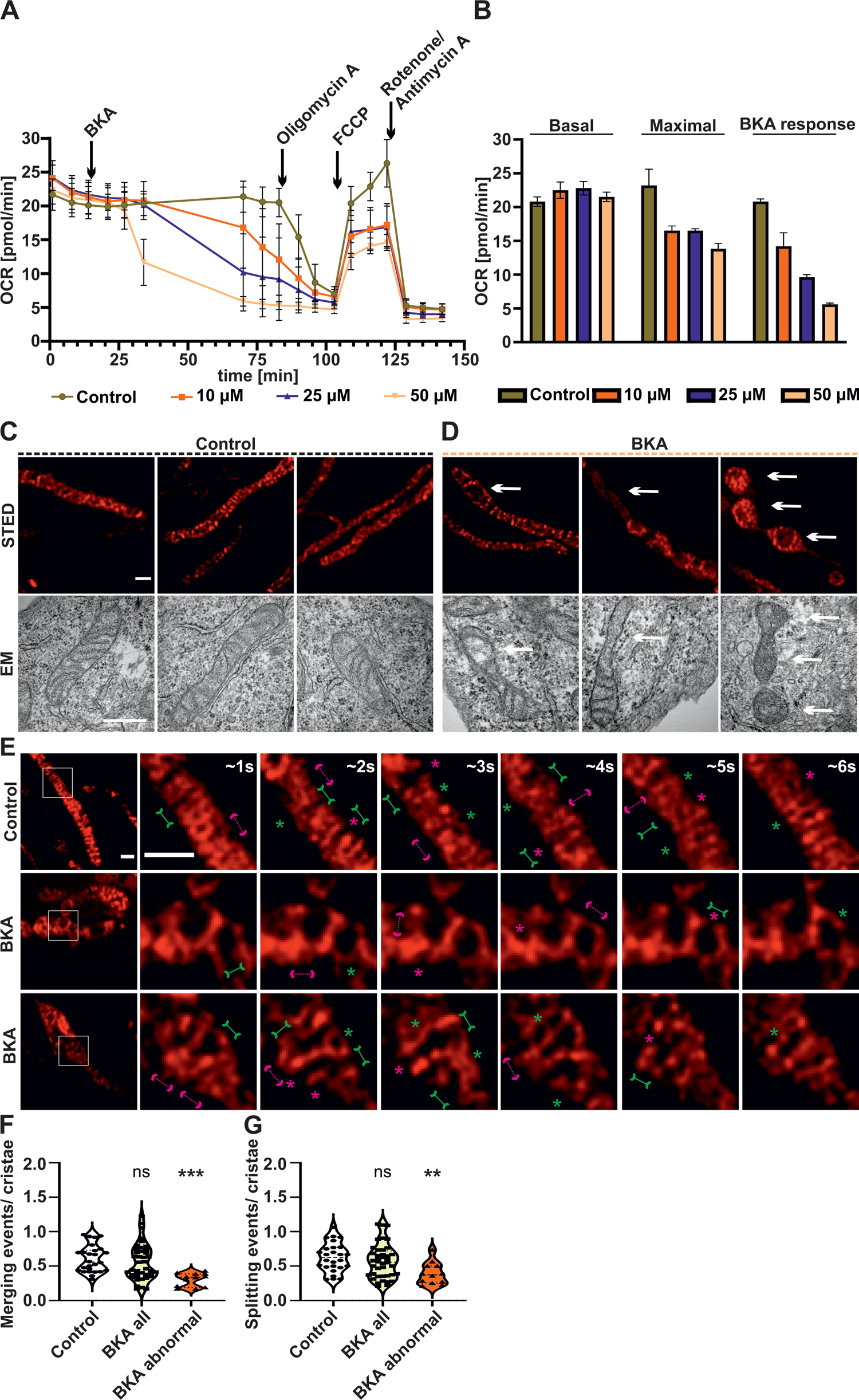
Inhibition of ANT causes perturbations in crista morphology and dynamics. **(A-B)** The oxygen consumption rates (pmol/min) of HeLa cells treated without or with various BKA concentrations (10 µM, 25 µM or 50 µM BKA, as indicated in color code) are shown. ∼70 min after BKA injection, routine Seahorse experiments measuring basal and maximal respiration were measured. Respective compound injection time-points are indicated by black arrows. Error bars represent standard deviation. **(B)** Detailed comparison of basal respiration (mean of first three measurements), maximal respiration (mean of three measurements after FCCP injection) and BKA response (mean of last three measurements before oligomycin A injection) of HeLa WT cells treated without or with various concentrations of BKA (10 µM, 25 µM or 50 µM BKA) as indicated using a color code. Error bars represent standard deviation. **(C-D)** Representative STED SR images (top row) of HeLa cells expressing ATP5I-SNAP, stained with silicon rhodamine treated without **(C)** or with **(D)** 50 µM BKA and corresponding electron micrographs of mitochondria (bottom row) displaying similar mitochondrial ultrastructure are shown. Three columns **(C)** display untreated mitochondria with normal morphology **(D)** Abnormal cristae morphology of BKA-treated mitochondria showing regions of sparse cristae. Similar perturbations in cristae morphology visualized by STED and EM images are indicated by arrows. Scale bars represent 500 nm. **(E)** Additional live-cell STED SR images of HeLa cells, from same conditions as **C, D** are shown. Images at the extreme left show whole mitochondria along with white inset boxes. Other images on the right-side display time-lapse series (0.94 s/frame) of zoom of mitochondrial portion at ∼1 s, 2 s, 3 s, 4 s, 5 s and 6 s. Green and magenta asterisks show corresponding merging and splitting events while solid green arrows pointing inward and dotted magenta arrows pointing outward show imminent merging and splitting events respectively. Scale bar represents 500 nm. **(F-G)** Blind quantification of cristae merging **(F)** and splitting **(G)** events per mitochondrion in HeLa cells treated without or with BKA. Mitochondria from BKA-treated cells were further separated into all mitochondria or those with exclusively abnormal cristae morphology and the individual groups compared to the untreated control (*ns = nonsignificant P-value* > 0.05, *\*\*P-value* ≤ 0.01, *\*\*\*P-value* ≤ 0.001). One-way ANOVA was used for statistical analysis.

## Discussion

The development of SR and high-resolution techniques which overcame the diffraction barrier of light, and their recent application to biological structures like mitochondria in fixed and living cells, has opened up exciting prospects to decipher mechanistic insights (Jakobs *et al*, 2020; Kondadi *et al*., 2020b). While EM could provide valuable insights into cristae remodelling by providing static data at different time-points, one could apply live-cell SR techniques like STED nanoscopy to understand the role of various proteins and metabolic factors regulating mitochondrial cristae dynamics. Here, we asked a basic question, namely whether modulation of OXPHOS, ΔΨ_m_, ATP levels, or ADP/ATP exchange in mitochondria determines cristae membrane dynamics, and if so, to which extent. In this study, we used advanced live-cell STED nanoscopy combined with newly developed and optimized quantification methods to study cristae morphology and dynamics when we inhibited the functioning of OXPHOS complexes I, III, V, and dissipated the ΔΨ_m_. Application of a set of well-characterized mitochondrial toxins led to formation of enlarged mitochondria, yet contrary to our expectations none of these toxins blocked cristae membrane dynamics. Before we discuss the details of the latter aspect, it is worth discussing the morphological alterations. Mitochondrial swelling is a phenomenon where there is an increase in the volume of the matrix caused due to osmotic imbalance between the matrix and cytosol (Kaasik *et al*, 2007). The osmotic balance is regulated by various channels and ion exchangers. Therefore, dysregulation of specific channels and exchangers in mitochondria could result in mitochondrial swelling. Additionally, opening of the mitochondrial permeability transition pore (MPT) causes mitochondrial swelling as the IM becomes permeable to solutes with a molecular weight less than 1.5 kDa (Lemasters *et al*, 2009). Mitochondrial swelling was proposed as mild reversible and excessive irreversible with the former regulating mitochondrial metabolism whereas the latter leading to mitochondrial dysfunction (Bernardi, 1999; Khmelinskii & Makarov, 2021a, b). The treatment of cells with mitochondrial toxins and imaging within a time-window of 30 min using live-cell STED nanoscopy suggests that the mitochondrial enlargement is in reversible mode with no loss of outer membrane which is consistent with our EM images. EM data from previous studies (Gottlieb *et al*., 2003; Hytti *et al*., 2019) are consistent with our live-cell STED nanoscopy and EM observations where the application of the described mitochondrial toxins led to structural alterations in enlarged mitochondria characterized by decreased cristae density. Consistent with our observations, it was shown that dissipation of ΔΨ_m_ by CCCP treatment led to decreased cristae density (Segawa *et al*, 2020). Concurrent to decreased cristae density, there was a trend of increased intercristae distance which was significantly higher in enlarged mitochondria after treatment with rotenone and CCCP. Overall, the cristae area was not changed when enlarged mitochondria were compared with normal mitochondria treated with mitochondrial toxins or not. Decreased cristae density along with no change in cristae area in enlarged mitochondria indicates presence of longer cristae compared to normal mitochondria in all treatment conditions.

Using live-cell respirometry and consistent with textbooks, it has been shown that mammalian cells instantaneously display decreased oxygen consumption upon inhibition of OXPHOS complexes I, III and V and increased oxygen consumption upon dissipation of ΔΨ_m_ using CCCP (Kondadi *et al*., 2020a; Stephan *et al*., 2020). Thus, addition of various mitochondrial toxins leads to opposing trends of oxygen consumption with CCCP displaying increased mitochondrial consumption as opposed to other three toxins. Unexpectedly, when the mitochondrial oxygen consumption was reduced after addition of rotenone, antimycin A and oligomycin A, we did not observe any change in the number of merging and splitting events in enlarged mitochondria when compared to normal mitochondria. On the contrary, increased oxygen consumption during CCCP exposure is connected to increased number of cristae merging as well as splitting events. Additionally, we demonstrated that the maintenance of the ΔΨ_m_ is not essential for cristae dynamics. Moreover, despite varying differences in cells treated with mitochondrial toxins w.r.t ΔΨ_m_, it can be concluded that largely no changes in the frequency of cristae dynamics were observed when the effects of different toxins are compared. Our data not only demonstrate that cristae membrane dynamics is not hampered upon loss of the membrane potential, it even shows an increase in merging and splitting events under these conditions. It should be noted that loss of ΔΨ_m_ is not a requirement for mitochondrial enlargement as cells treated with oligomycin A showed enlarged mitochondria but did not lose ΔΨ_m_. Overall, mitochondrial enlargement was necessary but not sufficient to display enhanced cristae membrane dynamics and these data point to the possibility that conditions of high oxygen consumption, which is equivalent to high electron flow from NADH to oxygen in the respiratory chain, promote cristae merging and splitting events.

What is the functional interplay between ATP levels and cristae dynamics? In order to decipher the ATP levels at the level of mitochondria, we used mitGoTeam2 probe which is a genetically-encoded sensor based on FRET for detecting differences in ATP levels (Nakano *et al*., 2011). The ATP levels are based on ratiometric FRET imaging meaning that the expression levels of the construct do not influence the ATP measurements. Further, it was shown that removal of glucose from the culture media results in a decrease of FRET ratio of mitochondrial ATP levels from 1.0 to ∼0.7 to 0.9 in various cell types (Depaoli *et al*, 2018). Consistent with this previous study, we found a decrease in ratiometric ATP levels of mitochondria. While we found a consistent decrease of mitochondrial ATP levels in all cells exposed to mitochondrial toxins (except CCCP), cells containing normal mitochondria already showed decreased ATP levels. Interestingly, there was no further decrease in mitochondrial ATP levels when compared to enlarged mitochondria indicating that the increased cristae dynamics served to maintain the already reduced ATP levels. It is interesting to note that the cristae merging and splitting was increased in enlarged mitochondria which coincided with maintenance of mitochondrial ATP levels upon CCCP treatment. Probably increased oxygen consumption and cristae dynamics regulate the ATP levels or vice-versa. Next, inhibition of the ANT translocator by BKA treatment led to increased percentage of mitochondria with abnormal cristae morphology. Overall, when analyzing all mitochondria at first, we did not observe significant changes in cristae membrane dynamics, yet we detected the subpopulations of mitochondria with apparent different cristae membrane dynamics. When we considered this and divided our population in mitochondria where cristae morphology was abnormal vs. normal, the cristae merging and splitting were significantly decreased compared to untreated mitochondria. This agrees with previous data which showed cristae dynamics were reduced in *MIC13* KO (Kondadi *et al*., 2020a). Mitochondrial ultrastructure was aberrant in *MIC13* KO due to loss of CJs. Therefore, we propose that cristae morphology and dynamics are interlinked. Given that enlarged mitochondria upon ΔΨ_m_ dissipation show enhanced cristae dynamics, we propose that cristae dynamics is possibly determined by structural constraints. In such a scenario highly, densely packed cristae would impose a constraint limiting cristae dynamics. It may also be that a reduction in cristae density is followed by an increased cristae fusion and fission rate serving as kind of a compensatory mechanism. Another aspect that appears to be important with respect to regulation of cristae membrane dynamics is the possible link to metabolic flux across the inner membrane. As discussed, we observe enhanced cristae membrane dynamics when the ΔΨ_m_ is dissipated resulting in increased oxygen consumption, a condition characterized by high electron and proton flux. In contrast, under conditions that hamper ADP/ATP exchange cristae membrane dynamics is partially blocked. Yet, neither isolated inhibition of OXPHOS complexes I, III, V, nor mild reduction in ATP levels hampered cristae membrane dynamics grossly. Future studies will have to dissect which metabolite fluxes are of particular importance and how they are interconnected. Yet, our study reveals important and partly unexpected insights into the interlink between different modes of OXPHOS modulation and cristae membrane dynamics.

## Materials and Methods

### Cell culture, transfection and mitochondrial toxin treatment

HeLa cells were maintained in DMEM cell culture media with 1 g/L glucose (PAN-Biotech), 1 mM sodium pyruvate (Gibco), 2 mM glutaMAX^TM^ (Gibco), Pen-Strep (PAN-Biotech, penicillin 100 units/ml and streptomycin 100 ug/ml) and 10% fetal bovine serum (PAN Biotech) at 37°C and 5% CO_2_. The cells were transfected with 1 µg of ATP5I-SNAP (Kondadi *et al*., 2020a) or 1 µg of mitGoTeam2 plasmid DNA using GeneJuice® (Novagen) reagent for 48 hours according to the manufacturer’s protocol. For live-cell SR imaging, HeLa cells expressing ATP5I-SNAP were stained with 3 µM SNAP-cell 647-SiR (Silicon rhodamine) (NEB) for 30 min, in FluoroBrite DMEM media (Gibco) without phenol red containing 10% fetal bovine serum (PAN Biotech), 1 mM sodium pyruvate (Gibco), 2 mM glutaMAX^TM^ (Gibco) and Pen-Strep (Sigma-Aldrich, penicillin 100 units/ml and streptomycin 100 ug/ml). After silicon rhodamine staining, cells were washed twice with FluoroBrite media. The third wash was done 10 min after the second wash after which mitochondrial toxins were added. Live-cell STED imaging was done in a time-window of 10-30 min after addition of toxins at 37°C and 5% CO_2_. The following concentrations of mitochondrial toxins (Merck) were used: rotenone (5 µM), antimycin A (10 µM), oligomycin A (5 µM), CCCP (10 µM) and BKA (10 µM, 25 µM or 50 µM).

### Live-cell STED super-resolution nanoscopy and quantification of cristae dynamics

Live-cell STED SR imaging was performed on Leica SP8 laser scanning microscope equipped with a 93X glycerol objective (N.A = 1.3) and a STED module. The samples were excited using a white light laser at 633 nm and the images were collected at emission wavelength from 640 nm to 730 nm using a hybrid detector (HyD) while using a pulse STED depletion laser beam at a wavelength of 775 nm. In order to increase the specificity of the signal, gating STED was used from 0.8 ns onwards. An optimised pixel size of 22 nm was used and images were obtained at a rate of 0.94 s/frame. Before every imaging session, the alignment of the excitation and depletion laser beams were optimised using 80 nm colloidal gold particles (BBI Solutions) in order to ensure the maximum possible resolution. Huygens Deconvolution software (21.10.0p0) was used to process the acquired images. The raw data images are provided. The STED videos were carefully analysed frame wise and manually quantified in a blind manner to account for the average number of merging and splitting events per cristae within a mitochondrion using ImageJ software (Fiji). The average number of merging and splitting events per mitochondrion was determined and the whole mitochondrial population belonging to a particular condition were represented using violin plots.

### Quantification of various parameters related to cristae morphology

As cellular bioenergetic status influences mitochondrial ultrastructure, we maximised the number of cells used for STED nanoscopy, where we quantified various cristae parameters including merging and splitting events, by considering only a maximum of two mitochondria from each cell when they were treated with or without toxins. We determined mitochondrial width, cristae density, average intercristae distance and percentage cristae area occupied by mitochondria using custom-made macros in Fiji. In order to determine the width of an individual mitochondrion in images obtained using STED nanoscopy, we used the average of three separate line scans covering the maximum diameter at both ends as well the centre of mitochondria which were roughly drawn equidistant from each other. Further, in order to determine the cristae density, a segmented line was manually drawn across the length of each mitochondrion and an intensity profile of the pixels across the length of the line was created. The algorithm detected the number of cristae by using the number of maximum intensity points of the graph along the length of mitochondrial plot profiles. The obtained cristae number was divided by the previously determined area of the respective mitochondrion to calculate the cristae number per µm^2^ which we termed cristae density.

To measure the average distance between cristae defined as intercristae distance (in nm), we used the previously acquired intensity profiles of mitochondria to determine the exact coordinates of each cristae in the image. Euclidean distances between cristae were calculated using a custom-made macro. Due to the drastic variations in mitochondrial and cristae morphology upon BKA treatment, the above described macro for determination of cristae number was not used. For these data sets, the number of cristae per mitochondrion was counted manually and used for normalization of merging and splitting events. Next, for calculation of the percentage cristae area occupied by mitochondria, we used a semi-automated batch processing custom-made macro. Cristae structures were manually selected by applying appropriate threshold on the images and the total mitochondrial area was selected by drawing the outline of the whole mitochondrion. The “Analyze Particles” function of Fiji was used to calculate the cristae area where structures less than five pixels were excluded. The macro divided the sum of all cristae area by the whole mitochondrial area and multiplied the result by 100 to acquire the percentage cristae area occupied by that particular mitochondrion.

### Electron microscopy

HeLa cells were grown in 15 cm petri dishes at 37°C with 5% CO_2_ and treated with respective mitochondrial toxins for 30 min which were then fixed with 3% glutaraldehyde, 0.1 M sodium cacodylate buffer at pH 7.2. Cell pellets were washed in fresh 0.1 M sodium cacodylate buffer at pH 7.2, before embedding in 3% low melting agarose. They were stained by incubating in 1% osmium tetroxide for 50 min followed by two washes for 10 min with 0.1 M sodium cacodylate buffer and one wash with 70% ethanol for 10 min. Samples are stained using 1% uranyl acetate/1% phosphotungstic acid mixture in 70% ethanol for 60 min. Graded ethanol series was used to dehydrate the specimen. The samples were embedded in spur epoxy resin for polymerization at 70°C for 24 hours. Ultrathin sections obtained using a microtome were imaged with a transmission electron microscope (Hitachi, H600) at 75 V which had a Bioscan 792 camera (Gatan).

### FRET-based microscopy to measure ATP levels

Cells expressing the genetically-encoded mitGoTeam2 were used to determine the ATP levels, kindly provided by Hiromi Imamura, Kyoto, Japan (Nakano *et al*., 2011). Single optical sections were obtained with a 93X glycerol objective (N.A = 1.3) using Leica SP8 confocal microscope maintained at 37°C and 5% CO_2_. The samples were excited at 471 nm and the green and orange emission channels were simultaneously obtained from 502 nm to 538 nm (termed 520 nm) and 568 nm to 592 nm (termed 580 nm) respectively as described (Nakano *et al*., 2011) in photon counting mode. In order to quantify the ratiometric images obtained, a semi-automated custom-made macro was designed using Fiji software to analyse the acquired images in a batch processing mode. The cells of interest were manually selected by drawing a region of interest (ROI). The obtained orange emission channel images (580 nm) were divided by respective green emission channel images (520 nm) by using the “Image Calculator” function of Fiji. A threshold was manually applied on the resulting ratiometric 32-bit float image in order to exclude background pixels using the “Clear Outside” command. In order to categorise cells containing either swollen or normal mitochondria, the cut off for swollen mitochondria was set to 650 nm in congruence with STED SR nanoscopy. If 85% of the mitochondrial population featured enlarged mitochondria, the cells were designated as swollen. Similarly, if 85% of the mitochondrial population featured mitochondria whose width was less than 650 nm, the cell was considered as having normal mitochondria. We measured the diameter of whole mitochondrial population in the respective cells using Leica Application Suite X software (version 3.7.1.21655).

### Determination of mitochondrial membrane potential (ΔΨ_m_)

HeLa cells were incubated with 20 nM TMRM (Invitrogen) and 50 nM MitoTracker^TM^ Green (Invitrogen) in DMEM cell culture media along with other supplements (mentioned above) for 30 min at 37°C followed by three washes. 10 min after addition of respective toxins, cells were imaged for 20 min in DMEM media containing 10 mM HEPES buffer (Gibco) and other supplements. Mitochondrial toxins were present in the media during imaging sessions. Imaging was done on spinning disc confocal microscope (PerkinElmer) using a 60x oil-immersion objective (N.A = 1.49). Single optical sections were obtained using excitation wavelengths of 488 nm (MitoTracker^TM^ Green) and 561 nm (TMRM). The microscope was equipped with a Hamamatsu C9100 camera. Image analysis including background subtraction and measurement of mean fluorescence intensity were performed using Fiji software after drawing ROI around individual cells.

### Live-cell respirometry

All the respiration measurements were performed using Seahorse XFe96 Analyzer (Agilent). HeLa cells were seeded in Seahorse XF96 cell culture plate (Agilent) at a density of 3.5 × 10^4^ cells per well overnight. Next day, cells were washed and incubated in basic DMEM media (103575-100; Agilent) supplemented with 10 mM glucose (Sigma Aldrich), 2 mM glutamine (Thermo Scientific), and 1 mM pyruvate (Gibco) at 37°C, with no CO_2_ incubation 1 h before the assay. Mitochondrial respiration was measured using Seahorse XF Cell Mito Stress Test kit (Agilent) according to the manufacturer’s instructions. Three concentrations of BKA (Sigma Aldrich) at 10 µM, 25 µM and 50 µM were tested. The dilutions of BKA and all Mito Stress Test components were prepared in Seahorse medium. Basal respiration was defined as initial oxygen consumption (pmol/min) based on three initial measurements prior to injection of BKA. The first three initial measurements to calculate basal respiration were done at 1.3, 7.8 and 14.2 min. The duration between two measurements is ∼6 min except when BKA was added where there was a gap of ∼30 min between 3 initial and final measurements. After this gap of 30 min after BKA addition, the last three measurements were averaged to calculate the BKA response. The measurements after BKA injection were followed by an incubation phase with oligomycin A, FCCP, rotenone and antimycin A in combination, standard compounds used in Seahorse live-cell respirometry experiments, where the average oxygen consumption was calculated from three values after addition of respective toxins. Maximal respiration was measured after addition of FCCP. Cell number was normalized after the run using Hoechst staining. Data were analysed using wave software (Agilent). Further calculations were done in Microsoft Excel and figure preparation in GraphPad Prism.

### Statistics and data representation

Statistical significance was tested by one-way ANOVA followed by Dunnett’s test for multiple comparisons against single control group or Šídák’s test for multiple comparisons of selected pairs with *ns = nonsignificant P-value* > 0.05, *\* P-value* ≤ 0.05, *\*\*P-value* ≤ 0.01, *\*\*\*P-value* ≤ 0.001, *\*\*\*\* P-value* ≤ 0.0001. For statistical analysis as well as data representation, GraphPad Prism (version 9.5.1) was used.

## Supplementary Information

The supplementary information consists of 4 figures and 13 movies in AVI format.

## Acknowledgements

We thank Andrea Borchardt and Tanja Portugal for excellent technical assistance in performing electron microscopy and molecular biological experiments. The STED SR and FRET-based ATP experiments were performed at the Centre for Advanced Imaging (CAi), HHU, Düsseldorf. We are grateful to Hiromi Imamura from Kyoto University, Japan for providing us with mitGO-Ateam2 plasmid used for detecting ATP levels.

## Author Contributions

M Golombek: Investigation, methodology, visualization, validation, data curation, formal analysis and writing–review and editing.

T Tsigaras: Visualization, investigation, formal analysis and software. Y Schaumkessel: Investigation and formal analysis.

S Hänsch: Validation, resources, methodology and software. S Weidtkamp-Peters: Resources and methodology.

R Anand: Supervision, validation, visualization, investigation and writing–review and editing.

AS Reichert: Conceptualization, supervision, project administration, writing–review and editing and funding acquisition.

AK Kondadi: Conceptualization, supervision, project administration, investigation, writing– original draft, review, editing and funding acquisition.

## Supplementary Figure legends

**Fig S1.**
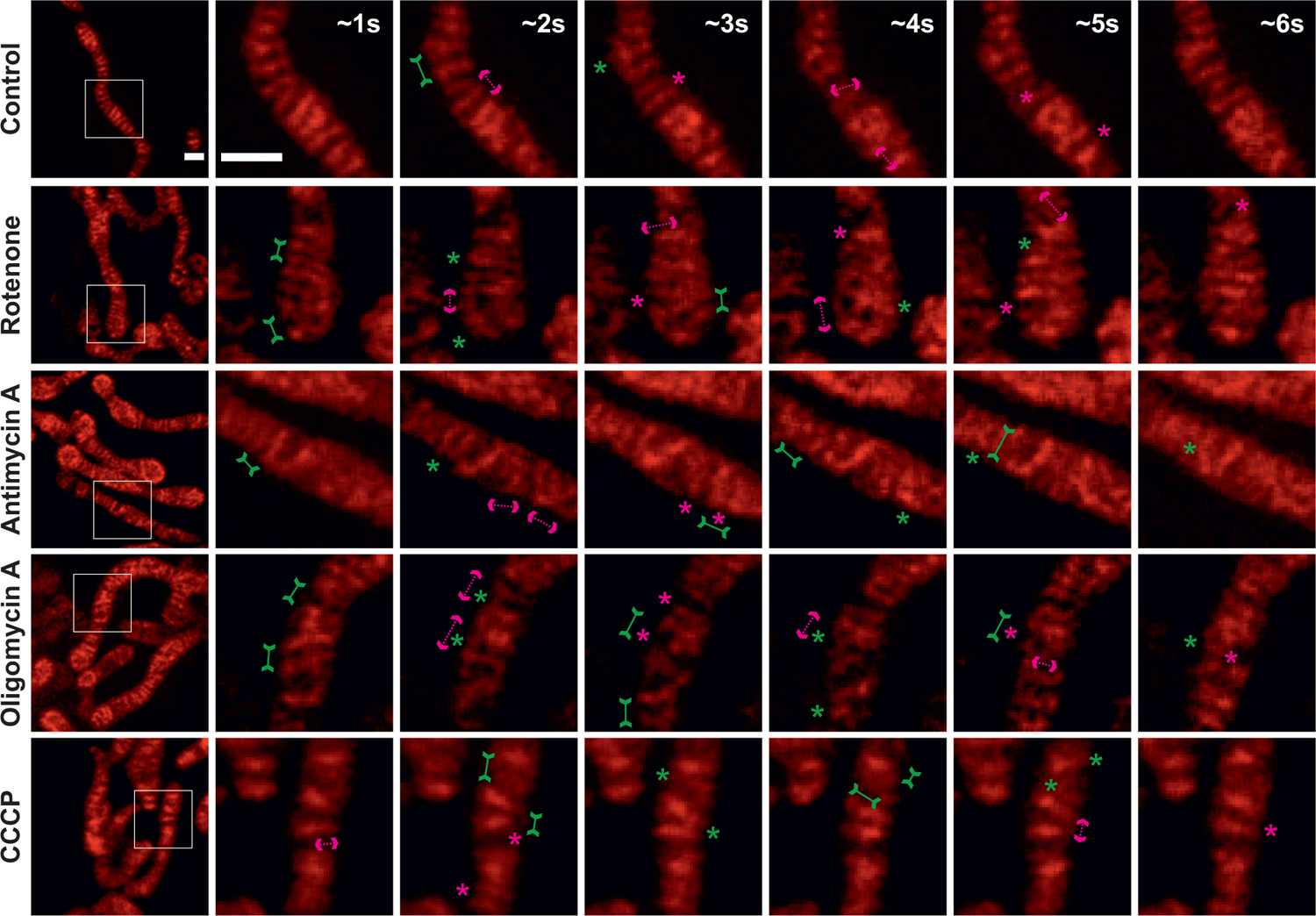
Crista merging and splitting events can be resolved in live-cell STED SR images even without deconvolution. Respective raw data of live-cell STED SR images, shown in **Fig 1A,** of HeLa cells expressing ATP5I-SNAP and stained with silicon rhodamine, untreated or treated with the rotenone, antimycin A, oligomycin A and CCCP. Images at the extreme left show whole mitochondria along with white inset boxes. Other images on the right-side display time-lapse series (0.94 s/frame) of zoom of mitochondrial portion at ∼1 s, 2 s, 3 s, 4 s, 5 s and 6 s. Green and magenta asterisks show corresponding merging and splitting events while solid green arrows pointing inward and dotted magenta arrows pointing outward show imminent merging and splitting events respectively. Scale bar represents 500 nm.

**Fig S2.**
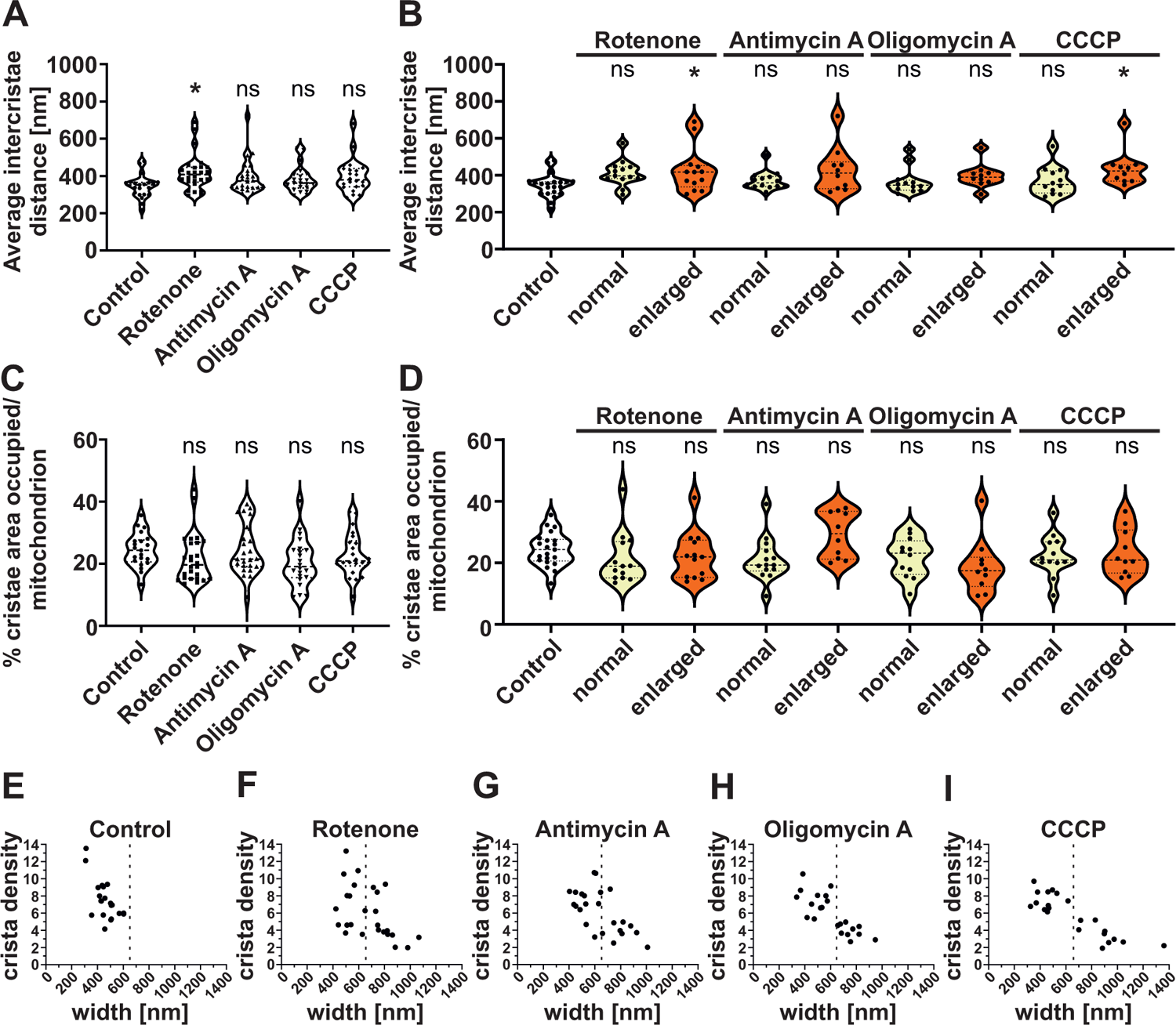
Mitochondrial enlargement correlates with reduced cristae density with no change in the cristae area. **(A-B)** Quantification of average intercristae distance (nm) per mitochondria, **(A)** Pooled data from three individual experiments is shown as violin plots with individual data points (21 to 26 mitochondria). Each symbol represents one mitochondrion. **(B)** Data was separated into normal (< 650 nm) or enlarged (≥ 650 nm) mitochondria with each condition having 10 to 21 mitochondria. Conditions are compared to untreated control group **(C-D)** Quantification of percentage crista area occupied per total mitochondrial area shown as violin plots with individual data points. Each symbol represents one mitochondrion. **(D)** Data was separated into normal and enlarged mitochondria (10 to 21 mitochondria). Conditions are compared to untreated control group. **(E-I)** Correlation of cristae density and mitochondrial width in control and toxin-treated conditions. Dotted line at 650 nm separates normal and enlarged mitochondria distributed in control cells **(E)** and cells treated with rotenone **(F)**, antimycin A **(G)**, oligomycin A **(H)** and CCCP **(I)** from three independent experiments with each condition including 21 to 26 mitochondria. (*ns = nonsignificant P-value* > 0.05, *\* P-value* ≤ 0.05). One-way ANOVA was used for statistical analysis.

**Fig S3.**
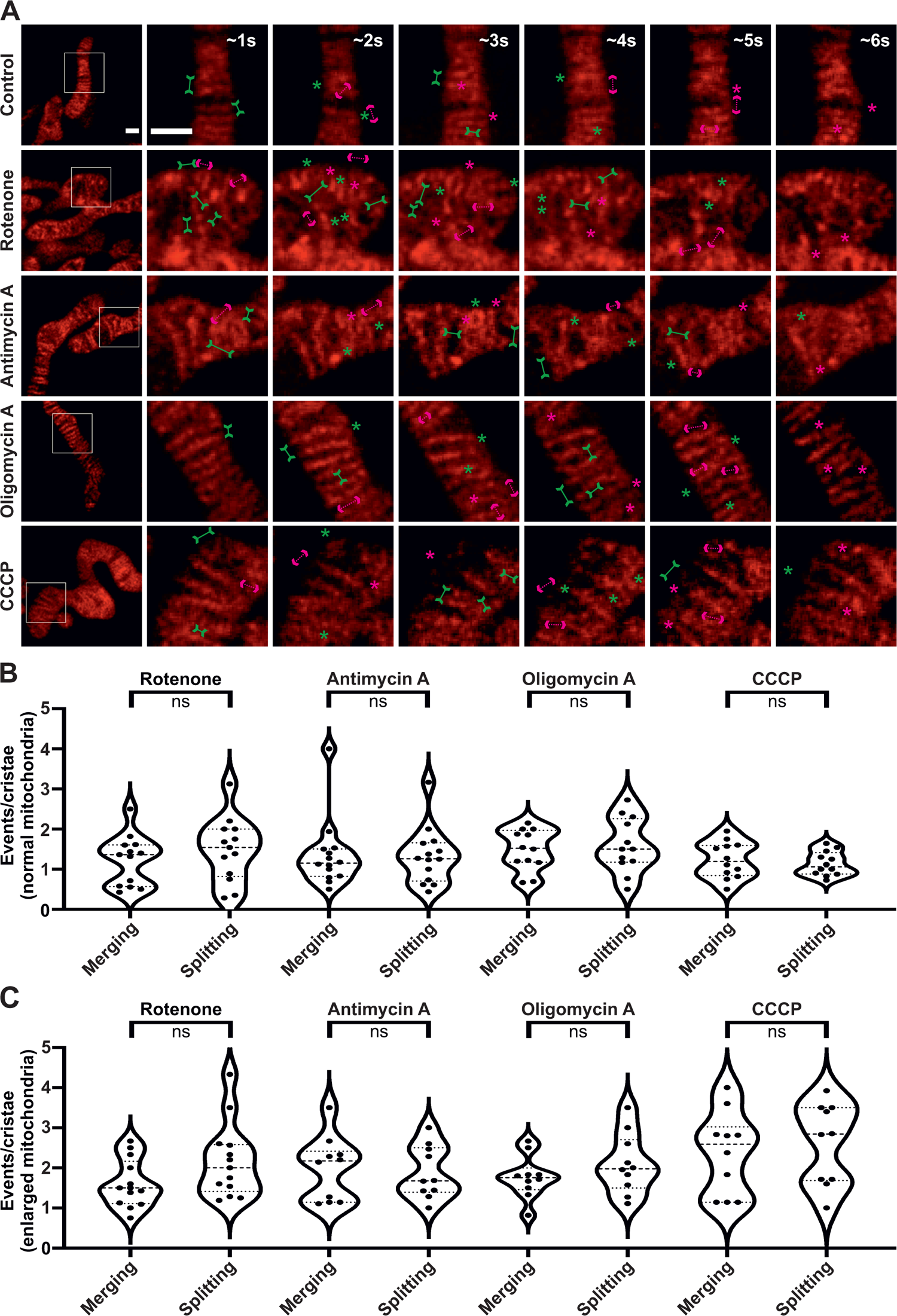
Crista merging and splitting events are present in a balanced manner in enlarged mitochondria. **(A)** Respective raw data of live-cell STED SR images shown in **Fig 3A** of HeLa cells, expressing ATP5I-SNAP and stained with silicon rhodamine, untreated (containing normal mitochondria) or treated (containing enlarged mitochondria) with the various mitochondrial toxins. Images at the extreme left show whole mitochondria along with white inset boxes. Other images on the right-side display time-lapse series (0.94 s/frame) of zoom of mitochondrial portion in the white inset at ∼1 s, 2 s, 3 s, 4 s, 5 s and 6 s. Green and magenta asterisks show corresponding merging and splitting events while green arrows pointing inward and dotted magenta arrows pointing outward show imminent merging and splitting events respectively. Scale bar represents 500 nm. **(B-C)** Blind quantification of merging and splitting events of cristae per mitochondrion in different conditions described in **(A)** and further separated into normal **(B)** and enlarged **(C)** mitochondria. Statistical analysis was performed within the individual treatment conditions using one-way ANOVA. (*ns = nonsignificant P-value* > 0.05).

**Fig S4.**
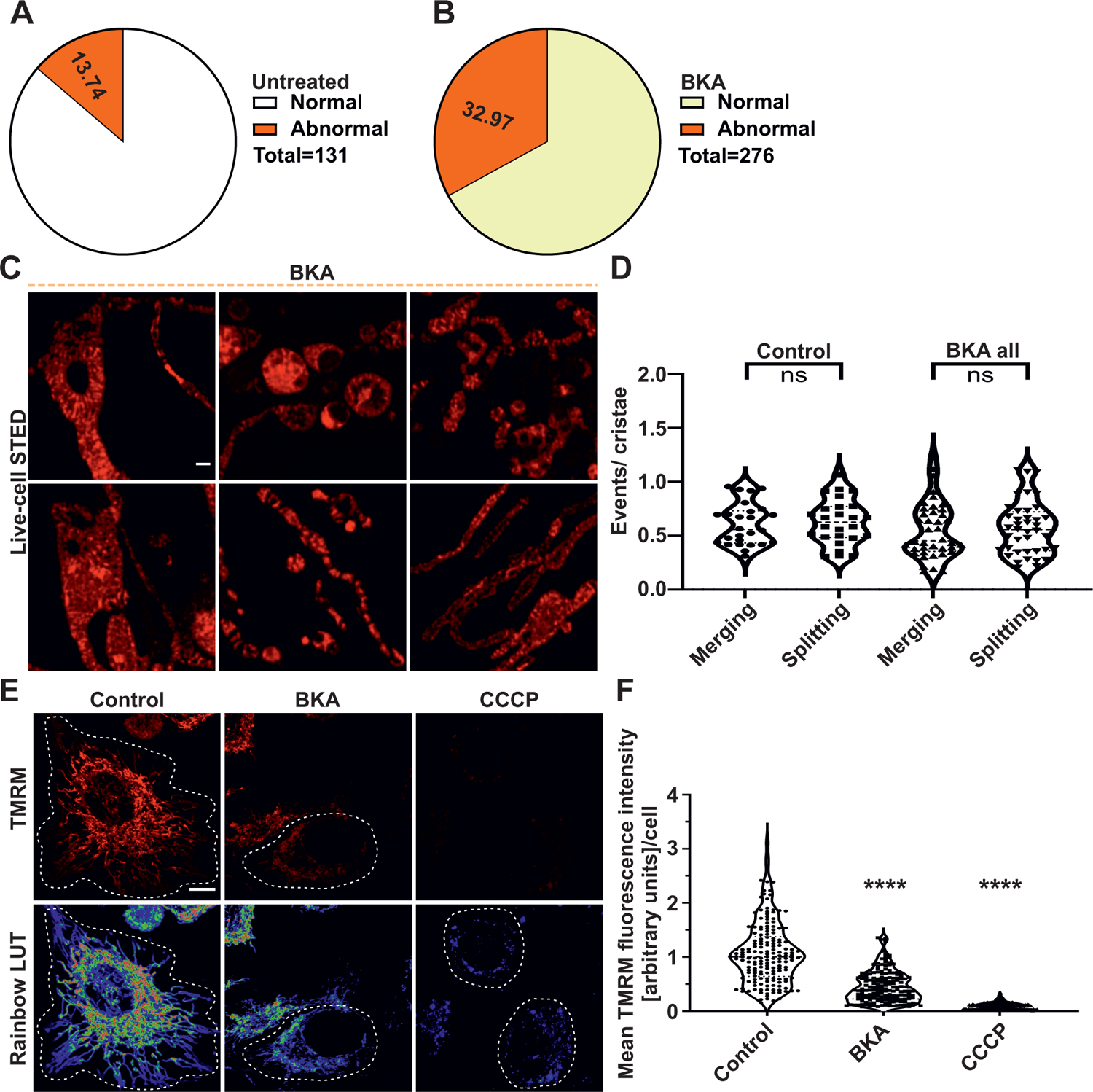
Inhibition of ANT perturbs cristae morphology and reduces ΔΨ_m_ without influencing the balance of merging and splitting events. **(A-B)** Pie charts showing percentage mitochondria with normal and abnormal cristae morphology in **(A)** untreated and **(B)** BKA-treated HeLa cells. 131 and 276 mitochondria from 54 and 95 STED SR images were considered respectively. **(C)** Representative STED SR images of HeLa cells expressing ATP5I-SNAP, stained with silicon rhodamine, display perturbed crista morphology in BKA-treated cells. Scale bar represents 500 nm. **(D)** Blind quantification of cristae merging and splitting events per mitochondrion when HeLa cells were treated with BKA or not. Pooled data from five separate experiments with 26 to 40 mitochondria is shown as violin plots with individual data points. Each symbol represents one mitochondrion. **(E)** Representative confocal images of control HeLa cells and cells treated with BKA or CCCP which were stained with TMRM are shown in top panel. Corresponding pseudocolor rainbow LUT intensities are shown in bottom panel. Scale bar represents 10 µm. **(F)** Quantification of ΔΨ_m_ based on TMRM mean fluorescence intensity measurements of individual HeLa cells that were not treated or treated with BKA or CCCP. Results are shown as violin plots including all individual data points. Data are obtained from 3 independent experiments, with each condition having 170 to 191 cells. Statistical comparisons were drawn between the untreated control group and the toxin-treated conditions. (*ns = nonsignificant P-value* > 0.05, *\*\*\*\* P-value* ≤ 0.0001). One-way ANOVA was used for statistical analysis.

## Supplementary Movie legends

**Movies 1 and 6:** Cristae merging and splitting events occur in a balanced manner at a time scale of seconds in untreated HeLa cells. Representative live-cell STED super-resolution movies (0.94 s/frame) of HeLa cell mitochondrion expressing ATP5I-SNAP, stained with silicon rhodamine, showing individual cristae and their dynamics. Time frame is indicated by a time stamper and the scale bar represents 500 nm. The white box highlights the region used to show individual merging and splitting events in the topmost row of Figs 1A and Fig 3A respectively.

**Movies 2, 3, 4 and 5** – Cristae merging and splitting is not hindered in normal mitochondria (< 650 nm) of HeLa cells treated with various mitochondrial toxins inhibiting complex I, complex III, complex V and dissipating ΔΨ_m_ respectively. Representative live-cell STED super-resolution movies (0.94 s/frame) of normal mitochondria (< 650 nm) treated with rotenone (**movie 2**), antimycin A (**movie 3**), oligomycin A (**movie 4**) and CCCP (**movie 5**). Mitochondria expressing ATP5I-SNAP, stained with silicon rhodamine, showing individual cristae reveal no hindrance in cristae dynamics. Time frame is indicated by a time stamper and the scale bar represents 500 nm. The white box highlights the region used to show individual merging and splitting events in Fig 1A.

**Movies 7, 8, 9 and 10** – Cristae merging and splitting is not reduced in enlarged mitochondria (≥ 650 nm) of HeLa cells treated with various mitochondrial toxins inhibiting complex I, complex III, complex V and dissipating ΔΨ_m_ respectively. Representative live-cell STED super-resolution movies (0.94 s/frame) of enlarged mitochondria (≥ 650 nm) treated with rotenone (**movie 7**), antimycin A (**movie 8**), oligomycin A (**movie 9**) and CCCP (**movie 10**). Mitochondria expressing ATP5I-SNAP, stained with silicon rhodamine, showing individual cristae reveal no reduction in cristae dynamics. Time frame is indicated by a time stamper and the scale bar represents 500 nm. The white box highlights the region used to show individual merging and splitting events in Fig 3A.

**Movie 11**: Cristae merging and splitting events occur in a balanced manner at a time scale of seconds in untreated HeLa cells (control for BKA-treated cells). Representative live-cell STED super-resolution movie (0.94 s/frame) of HeLa cell mitochondrion expressing ATP5I-SNAP, stained with silicon rhodamine, showing individual cristae and their dynamics. Time frame is indicated by a time stamper and the scale bar represents 500 nm. The white box highlights the region used to show individual merging and splitting events in the topmost row of Fig 6E.

**Movies 12 and 13**: Aberrant cristae structure accompanied by reduced cristae dynamics is observed in HeLa cells treated with ANT inhibitor, bongkrekic acid. Representative live-cell STED super-resolution movies (0.94 s/frame) of HeLa cells treated with BKA. Mitochondria expressing ATP5I-SNAP, stained with silicon rhodamine, show highly interconnected cristae with decreased merging and splitting events. Time frame is indicated by a time stamper and the scale bar represents 500 nm. The white box highlights the region used to show individual merging and splitting events in the two bottom rows of Fig 6E.

